# Structure of the receptor-activated human TRPC6 ion channel

**DOI:** 10.1101/282814

**Authors:** Qinglin Tang, Wenjun Guo, Li Zheng, Jing-Xiang Wu, Meng Liu, Xindi Zhou, Xiaolin Zhang, Lei Chen

**Affiliations:** State Key Laboratory of Membrane Biology, Institute of Molecular Medicine, Peking University, Beijing Key Laboratory of Cardiometabolic Molecular Medicine, Beijing 100871, China; Dizal Pharmaceutical Company, Jiangsu, China; Peking-Tsinghua Center for Life Sciences, Peking University, Beijing 100871, China

**Author notes:** These authors contribute equally to this work (Qinglin Tang, Wenjun Guo). Correspondence either to: Lei Chen, Or Xiaolin Zhang.

## Abstract

TRPC6 is a receptor-activated nonselective cation channel that belongs to the family of canonical transient receptor potential (TRPC) channels. It is activated by diacylglycerol, a lipid second messenger. TRPC6 is involved in many physiological processes and implicated in human genetic diseases. Here we present the structure of human TRPC6 homotetramer in complex with a newly identified high affinity inhibitor BTDM solved by single-particle cryo-electron microscopy to 3.8 Å resolution. The structure shows a two-layer architecture, in which the bell-shaped cytosolic layer holds the transmembrane layer. Extensive inter-subunit interactions of cytosolic domain, including N terminal ankyrin repeats and C terminal coiled-coil, contribute to the tetramer assembly. The high affinity inhibitor BTDM wedges between S5-S6 pore domain and voltage sensor-like domain to inhibit channel opening. Our structure uncovers the molecular architecture of TRPC channels and provides a structural basis for understanding the mechanism of these channels.

## Introduction

Studies on *Drosophila melanogaster* light sensing mutants reveal that the activation of rhodopsin by light results in currents mediated by TRP channel ^1,2^. Later, a few TRP channel families are identified in other species including mammals. These channels share sequence homology with the founding member drosophila TRP channel and response to various stimuli ^3^. Based on their primary sequences and activation mechanism, human TRPC3, TRPC6 and TRPC7 cluster on the phylogenic tree and constitute a receptor-activated TRPC subfamily. Similar to the drosophila TRP channel, they are non-selective cation channels that are directly activated by diacylglycerol (DAG) produced by receptor-coupled phospholipase C (PLC) ^4-6^. The activation of these channels results in the depolarization of cell membrane and calcium influx. TRPC3/6/7 can form either homo-tetramers or hetero-tetramers with variable calcium ion permeability ^7^. Based on their sequence features, these channels have a tetrameric transmembrane pore formed by six transmembrane helices, like canonical voltage-gated potassium channels. In addition, they have a large cytoplasmic N-terminus that contains four ankyrin repeats and a C-terminal coiled-coil motif. But how these structural elements are assembled into the tetrameric TRPC channel is largely unknown.

TRPC3/6/7 are broadly distributed in human tissues and involved in many physiological processes, such as synaptic formation ^8^, motor coordination^9^, kidney function ^10,11^, wounding healing ^12^, cancer progression ^13^, smooth muscle contraction ^14^, and cardiac hypertrophy induction ^15^. Notably, TRPC6 is important for kidney podocytes foot processes and genetic mutations in human TRPC6 can cause familial focal segmental glomerulosclerosis (FSGS)^10,11,16-18^, which is a kidney disease manifested by proteinuria^19^.

Despite the physiological importance of receptor-activated TRPC channels, their architecture and mechanism remain elusive due to lack of high resolution structures. To gain insights into the mechanism of receptor-activated TRPC channels, we embarked on the structural study of human TRPC6 channel and determined its structure in lipid nanodiscs. The structure reveals a new architecture of TRP channel family. Accompanied with functional data, we elucidated the structural basis for a novel high affinity inhibitor binding.

## Results

### Discovery of a high affinity inhibitor for hTRPC6

Gain of function mutations in human TRPC6 can lead to excessive calcium influx in kidney podocytes and familial FSGS ^20^. Currently, it is a clinical challenge to effectively cure this disease ^19,21-23^, and patients can easily develop end-stage renal disease, which can only be treated by frequent renal dialysis or kidney transplant. Pharmacological inhibition of TRPC6 is a promising way to alleviate clinical symptom in FSGS patients with TRPC6 gain of function mutations. Therefore, we screened small molecules that could inhibit hTRPC6-mediated membrane depolarization as lead compounds for the clinical treatment of FSGS. Through screening, we identified (2-(benzo[d][1,3]dioxol-5-ylamino)thiazol-4-yl)((3S,5R)-3,5-dimethylpiperidin-1-yl)methanone (abbreviated, BTDM) as a high-affinity inhibitor for the hTRPC6 channel (Fig. 1a). We used 1-oleoyl-2-acetyl-sn-glycerol (OAG), a soluble DAG analogue to activate hTRPC6 channel. We found BTDM potently inhibited OAG-evoked currents and membrane depolarization of cells transfected with the hTRPC6 expression constructs (Fig. 1b-d). In addition, BTDM could suppress the activity of gain of function hTRPC6 channel mutants, including P112Q ^11^, R895C and E897K ^10^, with similar potency as the wild-type channel (Supplementary information, Table S1), further emphasizing the potential of BTDM as a lead compound.

**Figure 1.**
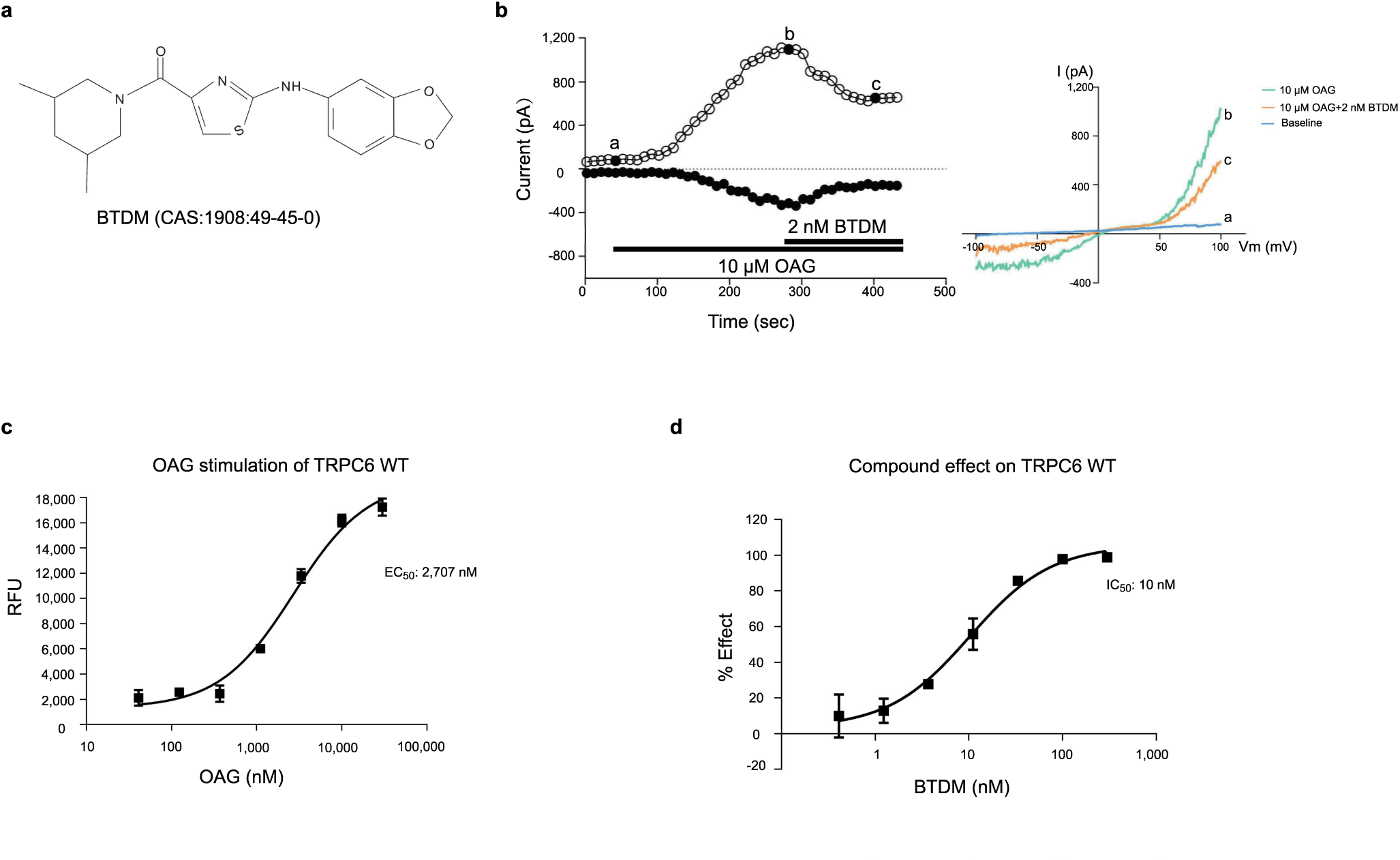
Characterization of the high affinity inhibitor BTDM. **a,** Chemical structure of the BTDM used in this study. **b,** Whole cell currents of hTRPC6 evoked by OAG were inhibited by BTDM in sub-IC50 concentration. Voltage ramps were applied every 10 s, and the outward current at +100 mV and inward current at −100 mV during each ramp are plotted as a function of time after rupture of the patch for whole cell recording. Solution exchanges were indicated by horizontal bars. I-V curves were obtained during voltage ramps at the time points indicated. The inhibitor could inhibit hTRPC6 current with the inhibition rate of 45.54% ± 2.22 at 2 nM (n=3). **c,** Activation effect of OAG on hTRPC6 constructs, measured by FLIPR membrane potential assay (n=3). **d,** Inhibition effect of the BTDM on wild type hTRPC6 (n=3).

### Architecture of the hTRPC6 channel

To stabilize hTRPC6 in the closed state, we conducted cryo-electron microscopy (cryo-EM) studies of hTRPC6 in complex with the inhibitor BTDM. hTRPC6 protein overexpressed in mammalian cells was purified in detergent micelles and reconstituted into nanodiscs for single particle analysis (Supplementary information, Figures S1 and S2). The 3.8 Å cryo-EM map was of sufficient quality for us to model 670 out of 931 residues (Supplementary information, Figure S1 and S2 and Table S2). The remaining residues were mostly disordered, probably due to their flexibility in the closed state. The hTRPC6 channel tetramer occupies 75 Å × 75 Å × 150 Å in the three-dimensional space (Fig. 2 and Supplementary information, Movie S1). Four protomers pack symmetrically to generate a two-layer architecture: the intracellular cytoplasmic domain (ICD) layer, and the transmembrane domain (TMD) layer (Fig. 2b,c). The central four-fold rotation axis is perpendicular to both layers. The hTRPC6 ICD has an inverted bell shape and caps below the ion channel pore of the TMD. Its N-terminus (1–84) was unresolved in the cryo-EM density map. Truncation of the N-terminal amino acids 1–71 did not affect tetramer assembly or OAG-induced depolarization of membrane potential (Supplementary information, Figure S5 and Table S1), indicating that the flexible N terminal 71 residues were dispensable for hTRPC6 assembly and gating. Four ankyrin repeats (ARs, residues: 96–243) and adjacent linker helices (LHs, residues: 256–393) form the major building block of ICD (Fig. 3). The last three LHs pack against the TRP helix, providing the major contact site between the ICD and TMD (Fig. 3b,c). Immediately after the TRP helix, the TRP re-entrant dips halfway into the phospholipid bilayer (Fig. 3b,c). The amino acids of the cytoplasmic C-terminus of TRPC6 fold into two long helices, namely C-terminal helix 1 (CH1) and C-terminal helix 2 (CH2). CH1 runs horizontally from the periphery into the center of the channel and connects to the vertical CH2 coiled-coil via a 90°turn (Fig. 3). The amino acids between the TRP re-entrant and CH1 (766– 854) are disordered. These residues contain putative binding sites for inositol hexaphosphate (IP_6_) (842–868) ^24^ and the calmodulin and IP_3_ receptor, namely the calmodulin/IP_3_ receptor binding (CIRB) region (838–872) ^24-26^. It is reported that calmodulin and IP3 receptor can bind to ICD of TRPC channels to modulate their activity ^24-26^.

**Figure 2.**
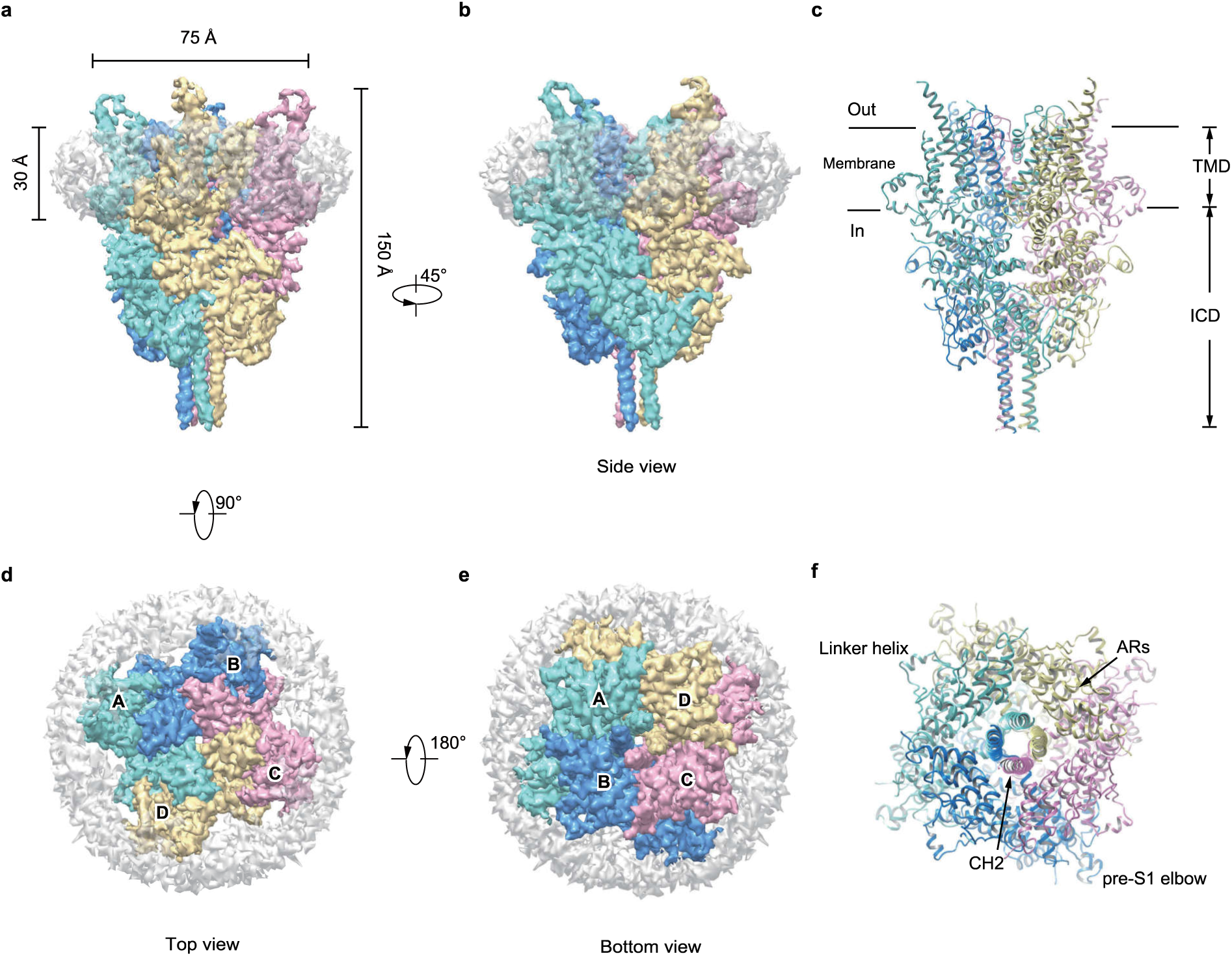
Cryo-EM density map of the human TRPC6 channel. **a, b,** Cryo-EM map of hTRPC6 channel is shown in side view. Subunits A, B, C and D are colored in cyan, blue, pink and yellow, respectively. Density corresponding to nanodisc is in grey with transparency. c, hTRPC6 model is shown in cartoon from side view. **d, e,** Top view (**d**) and bottom view (**e**) of the hTRPC6 channel density map. f, Bottom view of hTRPC6 model. AR, ankyrin repeat; CH, C-terminal helix.

**Figure 3.**
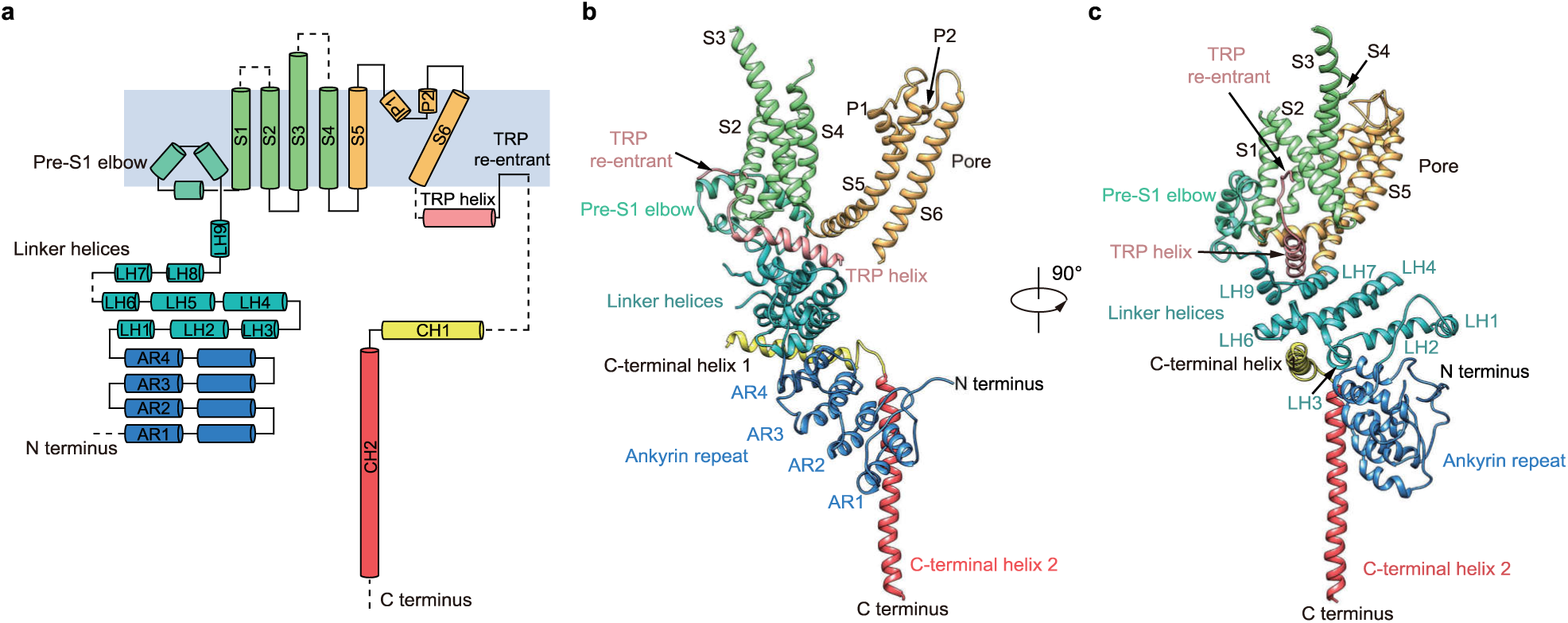
Structure of a single TRPC6 subunit. **a,** Topology and domain organization of hTRPC6. α-helices are shown as cylinders. Dashed lines represent flexible regions with insufficient density for model building. **b, c,** Ribbon diagrams show two views of one TRPC6 subunit with the same color as in **a**.

### Inter-subunit interactions mediate hTRPC6 tetramer assembly

Both the ICD and TMD contribute to the tetrameric assembly of the hTRPC6 channel. The ICD of each subunit extensively interacts with the neighboring subunits, forming an inter-subunit interface with the area of 5,483 Å^2^ between adjacent subunits (Fig. 4). Specifically, the amino acids of the N-terminal loop (85–94) interact with the LHs of the neighboring subunit (Fig. 4d). The LHs also interact with adjacent LHs (Fig. 4e). Similar interactions were previously mapped biochemically in TRPC4, in which N-terminal 1–304 residues interact with both residues 87–172 and residues 254–304 ^27^. Moreover, CH1 inserts into the cavity between two neighboring subunits and glues them together (Fig. 4f,h). CH2 helices form a vertical four-helix bundle that further tightens the tetramer (Fig. 4g), reminiscent of TRPA1 and TRPM C-terminal four-helix bundle structure ^28-32^. Uniquely, the four CH1–CH2 connection linkers pack tightly to form a knot-like structure on the top of CH2 (Fig. 4i). Deletion of the C-terminal half of CH2 (905–931) did not dramatically affect hTRPC6 assembly and gating (Supplementary information, Figure S5 and Table S1) but further deletion of entire CH2 and CH1–CH2 linker (878–931) greatly impaired tetrameric channel assembly, as shown by dominant dimer peak on fluorescence-detection size-exclusion chromatography (FSEC) (Supplementary information, Figure S5a), indicating the essential role of the CH1–CH2 linker knot and the first half of the CH2 in tetramer assembly.

**Figure 4.**
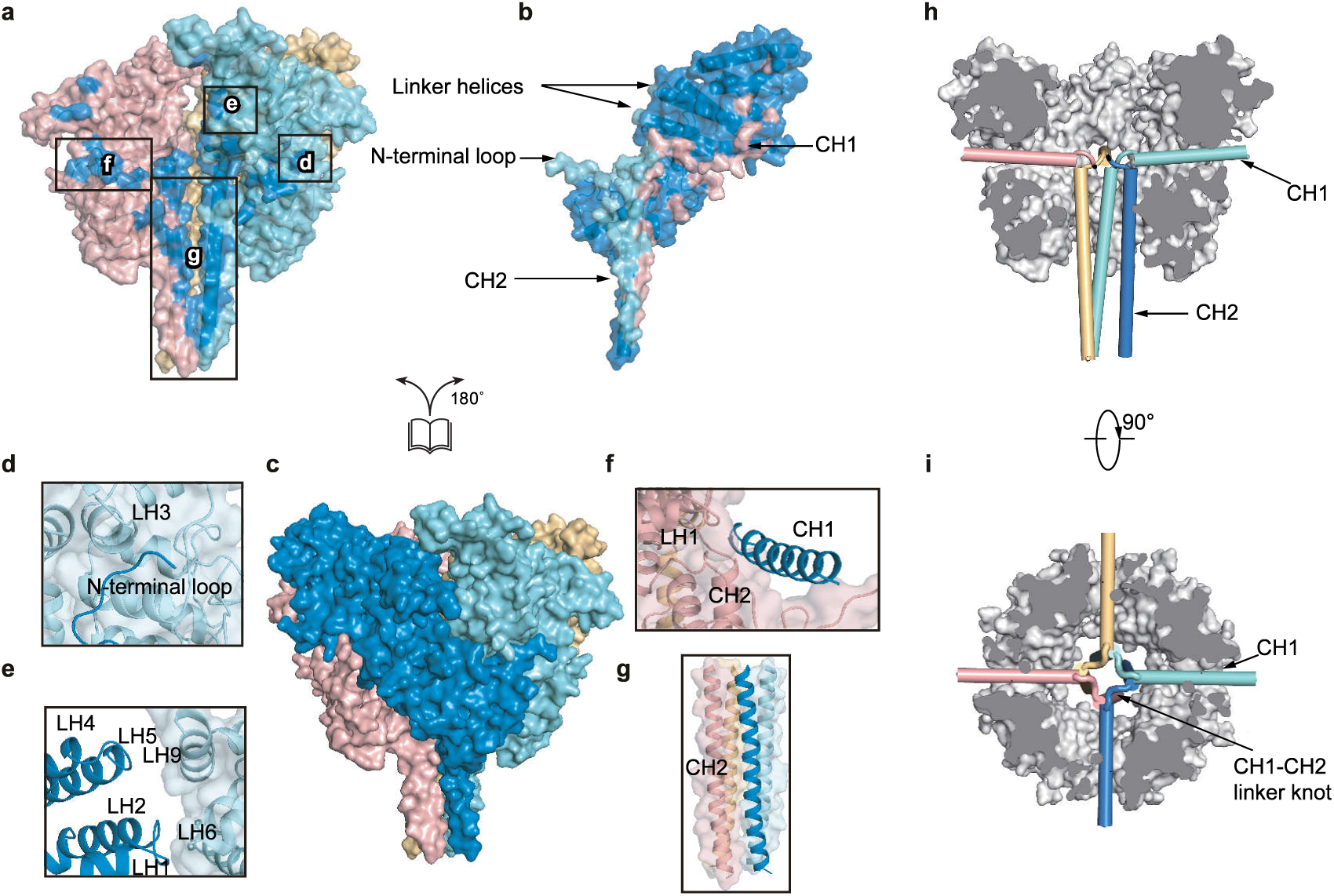
Inter-subunit interface of hTRPC6 ICD. **a-c,** Open-book view of hTRPC6 ICD inter-subunit interface shown in surface representation. Surface of each subunit is in transparency and colored the same as in Figure 1. View of the ICDs of subunits A, C and D is shown in **a** and residues that interact with the subunit B are colored in blue. View of the ICD of subunit B is shown in **b** and residues that interact with subunits A, C and D are colored in cyan, pink and yellow, respectively. View of the ICD layer tetramer is shown in **c**. d-g, Close up view of the inter-subunit interactions boxed in **a**. **h,** Side view of CH1 and CH2 locations. CH1 and CH2 are shown as cylinders and colored the same as in **c**. The other parts of ICD are colored in grey. CH2 of subunit B is omitted for better illustration. **i,** Top view of CH1 and CH2 locations.

### Transmembrane domain of the hTRPC6 channel

In the TMD, the voltage sensor-like domains (VSLD, S1–S4) dock onto the 4-fold symmetric ion channel pore (S5–S6) in a domain-swapped fashion, as evidenced by the strong density of S4-S5 linker (Fig. 5 and Supplementary information, Figure S3d). This is similar to the conventional voltage-gated potassium channel ^33^ and other TRP channels ^34-42^. It is worth noting that the S3 helix of hTRPC6 is longer than that of other TRP channels for about 3 turns in the extracellular C-terminus. Together with extracellular loops that connect transmembrane helices, the S3 C-terminus creates a distinctive large protrusion from TMD into the extracellular space (Fig. 2a-c). These protrusions might harbor the extracellular calcium sites that are important for the channel activity regulation ^43^. Among all of the available TRP channel structures, the TMD of hTRPC6 mostly resembles that of NOMPC ^39^ and TRPM ^29-32^, as they share common structural elements of the pre-S1 elbow and the TRP re-entrant located after the TRP helix. According to the sequence alignments, these structural elements are also preserved in the drosophila TRP channel, and other mammalian TRPC channels (Supplementary information, Figure S4). The calculated pore profile showed that the radius of the narrowest constriction is 0.96 Å which is smaller than that of dehydrated sodium ions (Fig. 5c). Therefore, the pore in the hTRPC6 structure is in a closed state. This is consistent with the fact that the high-affinity inhibitor BTDM was supplemented in our cryo-EM sample preparation. The polar residues N278 and Q732 and hydrophobic residue I724 line the narrowest constriction and form the gate on the intracellular side (Fig. 5c). On the extracellular side, the negatively charged E687 sits on top of the selectivity filter (Fig. 5c-e). It has been reported that the same residue on TRPC3 (E630 of TRPC3) governs divalent permeability ^44^, consistent with its crucial position in our structure model.

**Figure 5.**
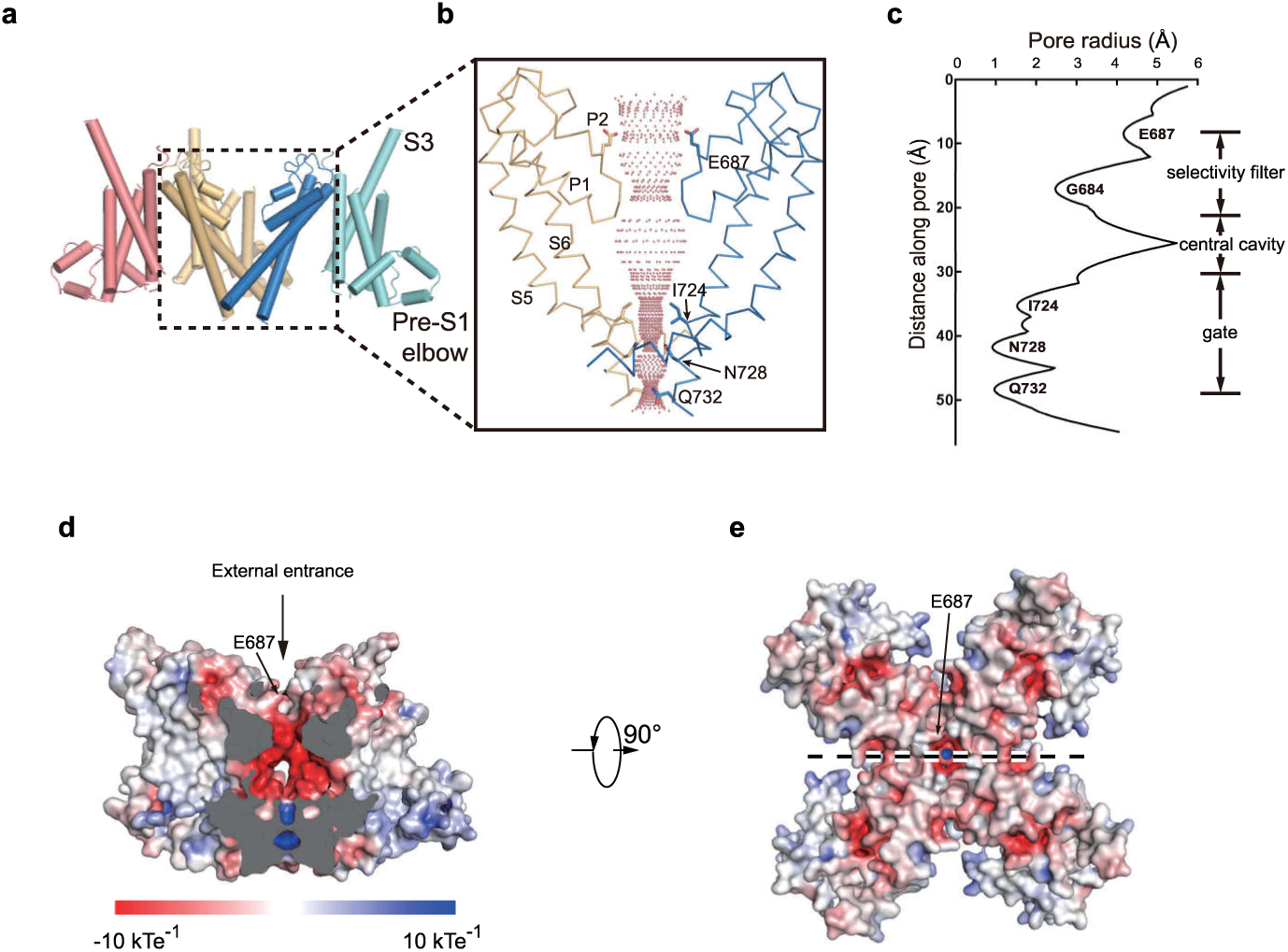
Structure of the hTRPC6 TMD. **a,** Side view of the hTPRC6 TMD in cartoon representation. S5–S6 of the subunits B and D and S1–S4 of the subunit C are omitted for better illustration. **b,** Close-up view of the S5–S6 pore region boxed in **a**. Ion-permeation pathway along the pore was calculated with the HOLE program. **c,** Calculated pore radius corresponding to **b** is shown vertically. **d,** Central cross-section of hTPRC6 transmembrane domain shown as surface and colored by electrostatic potential. Arrow indicates the external ion entrance. Position of E687 on top of the selectivity filter is indicated by an arrow. **e,** Top view of hTPRC6 transmembrane domain. A dashed line in **e** indicates the position of central cross-section in **d**.

### TRP domain of the hTRPC6 channel

TRP helix is sandwiched between the cytosolic LHs and the TMD (Fig. 6a). The LH7, LH8 and LH9 form a hydrophobic pocket to accommodate the bulky side chain of F744, W750, Y751, and F754 on the cytosolic face of TRP helix (Fig. 6b). On the membrane face of TRP helix, W742, L749 and Y753 tightly pack against S1, S4 and S4-S5 linker of TMD (Fig. 6c). TRP re-entrant interacts with both preS1 elbow and S1 helix (Fig. 6d). Similar reentering loop was also observed in the structures of NOMPC channel ^39^ and TRPM channel ^29-32^. It has been reported that the conserved site ^755^EE^756^ located between TRP helix and TRP re-entrant plays an important role in the STIM1-mediated gating of TRPC channels ^45^, correlated with their crucial position (Fig. 6d).

**Figure 6.**
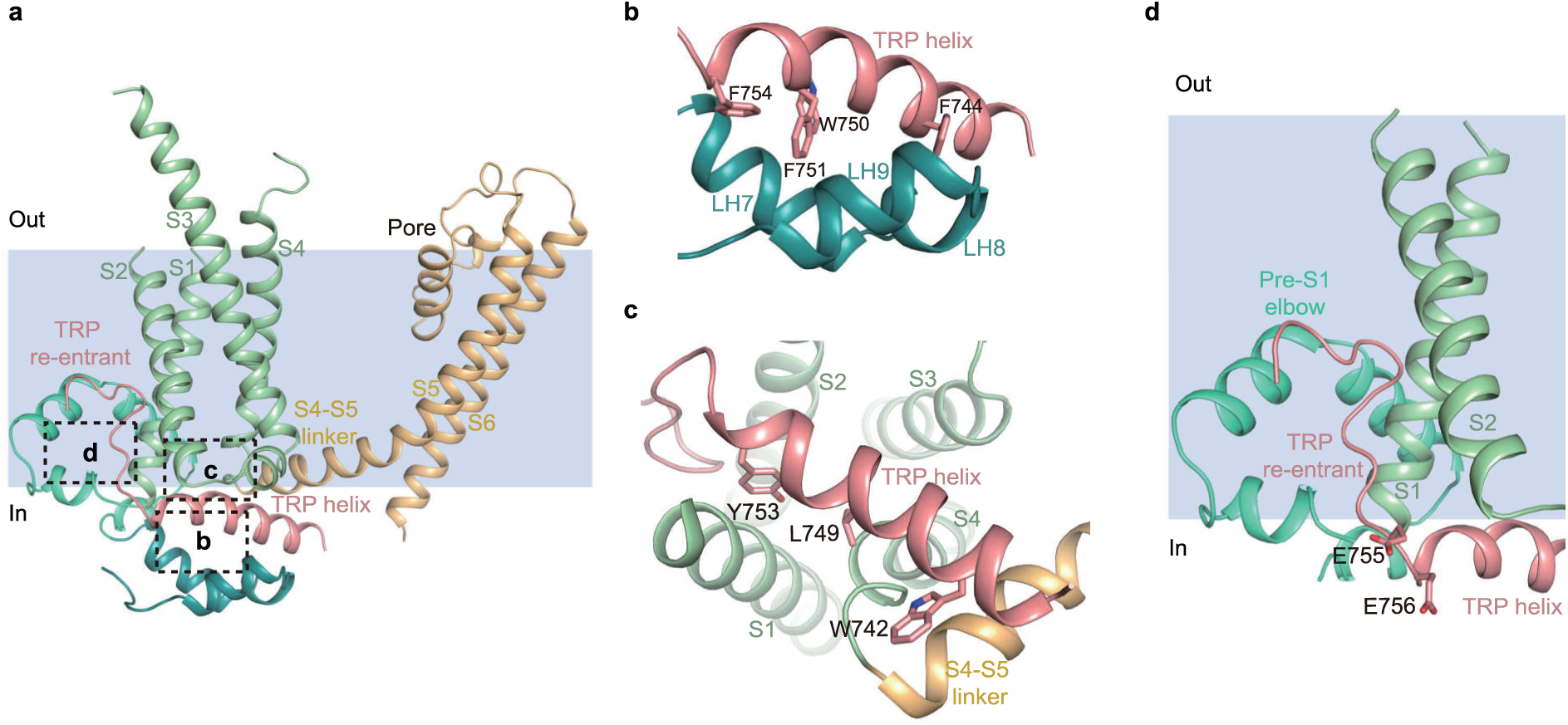
TRP domain of hTRPC6. **a,** Side view of LHs and TMD of one hTRPC6 subunit. Each structure element is colored the same as in Figure 2. Position of putative membrane bilayer is shown in light blue. **b,** Close-up view of interactions between TRP helix and LHs, boxed in **a**. **c,** Bottom view of the interactions between TRP helix and VSLD, boxed in **a**. **d,** Close-up view of the TRP re-entrant boxed in **a**.

### The inhibitor BTDM binding site

We found two extra densities in the map of the hTRPC6 TMD. One elongated extra density is surrounded by the pre-S1 elbow, S1, S4 and S4–S5 linker from one subunit and S5 from the adjacent subunit (Supplementary information, Figure S6a). To determine if the surrounding residues might be involved in BTDM binding, we made mutations of Y392F on the linker helix, A404C, F407A, L408M and L411F on the pre-S1elbow; A447T, A447F, T451F and T451L on S1; N617S and F620L on the S4-S5 linker (Supplementary information, Figure S6a). These mutations did not markedly affect OAG activation and BTDM efficacy (Supplementary information, Table S1). Notably, in the TRPM4 structure ^46^, a CHS molecule occupies the similar position (Supplementary information, Figure S6b). Therefore, we speculate that this density corresponds to a CHS molecule used in protein purification. The other extra density in the hTRPC6 TMD is surrounded by S3, S4 and S4-S5 linker of one subunit and S5-S6 of the adjacent subunit (Fig. 7 and Supplementary information, Figure S6c). The position of this density is similar to the resiniferatoxin (RTX) and the capsaicin-binding site on TRPV1 ^47^ (Supplementary fig. 6d). Residues W526 on S3, S608 on S4, Q624 and L627 on the S4-S5 linker from one subunit, together with residues I640 and V644 on S5, and T714 on S6 from the adjacent subunit create the binding pocket for the ligand represented by the extra density (Fig. 7b,c). We examined whether these residues are involved in BTDM binding by mutagenesis. We found that W526A and I640F had modest effects on BTDM function, whereas T714F and Q624A dramatically reduced BTDM affinity (Supplementary information, Table S1). Strikingly, BTDM had little effect on S608L and V644F mutants, even at 10 µM concentration (Supplementary information, Table S1). These mutants had normal OAG activation but impaired BTDM inhibition (Supplementary information, Table S1), emphasizing the importance of these residues on BTDM binding and suggesting that this pocket with extra density bound inside might represent the BTDM binding site on hTRPC6 channel. However, BTDM is a molecule with many possible rotamer conformations and limited local map quality precluded accurate modeling of BTDM into the extra density. Thus the detailed binding mode of BTDM needs further investigation.

**Figure 7.**
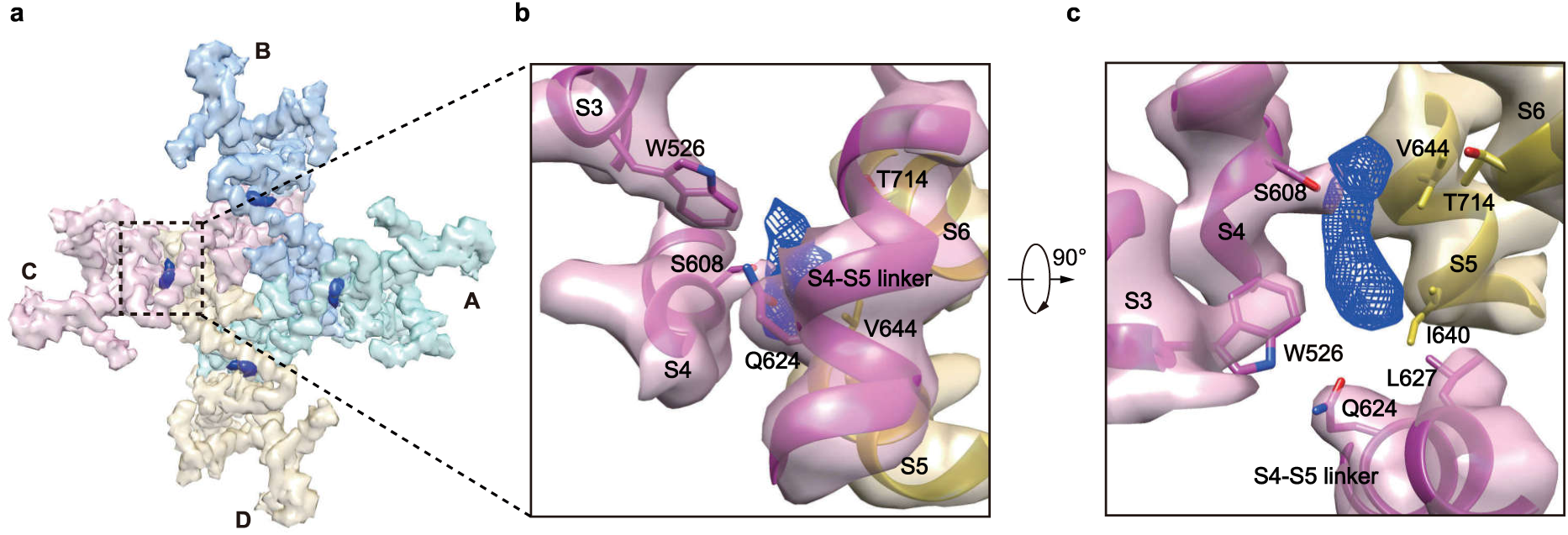
BTDM densities observed in hTRPC6 structure. **a,** Cryo-EM density map of hTRPC6 transmembrane domain in bottom view. Each subunit is colored the same as in Figure 1. Protein density is in transparency. BTDM density is colored in blue. **b, c,** Close-up view of BTDM binding site from bottom and side views.

## Discussion

The structures of hTRPC6 presented here provide a view of the architecture of this receptor-activated TRPC channel family. The large ICD harbors important sites for modulatory protein binding ^24-26^ and upstream kinase phosphorylation ^49^. It acts as an antenna to sense these intracellular signals and talks to the TMD via the TRP domain that bridges ICD and TMD (Supplementary information, Figure S7). The TRPC TMD harbors not only the cation permeation pathway but also the site for ligand binding. High affinity inhibitor BTDM wedges between the VSLD and the pore to inhibit channel gating. Our structures not only provide a template for structure-based drug discovery for hTRPC6 but also pave the way to understand the gating mechanism of receptor-activated TRPC family channels.

## Materials and methods

### Cell Culture

Sf9 cells were cultured in Sf-900 III SFM medium (Thermo Fisher Scientific) at 27 °C. Free style 293F cells were cultured in Freestyle 293 medium (Thermo Fisher Scientific) supplemented with 1% FBS at 37 °C with 6% CO_2_ and 70% humidity.

### Electrophysiology

hTRPC6 channel activation was measured using the whole-cell patch-clamp technique. 10 µM OAG was used to activate current in hTRPC6 stable expressing cells (TRPC6-HEK293, PerkinElmer Company). Cells were placed in a small chamber and continuously perfused with an external solution (∼3 ml/min) using a RSC-200 perfusion system (Science Instruments, BioLogic). All current recordings were conducted at room temperature (∼22°C). Electrodes were made from glass capillary tubes and had a resistance of 2-4 ΜΩ when filled with one of the internal solutions. For hTRPC6 current recording, the intracellular solution contains (in mM): 120 CsOH, 120 Aspartate, 20 CsCl, 2 MgCl_2_, 0.4 CaCl_2_, 10 HEPES, 2 Na_2_ATP, 0.1 Na_3_GTP, 10 Glucose and 1 EGTA (pH7.2-7.25, adjusted with CsOH). The extracellular solution contains (in mM): 145 NaCl, 5 KCl, 1 CaCl_2_, 1 MgCl_2_, 10 HEPES and 10 Glucose (pH 7.4 adjusted with NaOH). After whole-cell configuration was established, cell membrane capacitance was canceled electronically and the series resistance was compensated by about 70%. TRPC6 current was induced by a 300 ms voltage ramp protocol (from +100 mV to −100 mV) every 10 seconds at a holding potential of −60 mV. Once the control current is stabilized, the recording chamber is perfused with external solution containing testing compound. The TRPC6 current is measured as the average current at +100 mV. The time course of current is plotted for the whole experiment. Percent inhibition = 100 × (1 - l_D_/l_c_), where l_D_ is the current amplitude measured at the end of a particular drug concentration and l_c_ is the control current amplitude measured before drug application. Zero current (background) level was set at the very beginning before OAG activated TRPC6 current. MultiClamp 700B amplifier with Digidata 1440 interface and pCLAMP software (AXON Instruments) were used for data acquisition and analysis. All the reagents are purchased from Sigma Company.

### FSEC

The cDNA of hTRPC6 was cloned into a home-made N-terminal GFP-tagged BacMam vector ^50^. Various hTRPC6 constructs were screened by transfecting vectors into FreeStyle 293-F cells (Thermo Fisher Scientific, grown in 293Ti medium, 37°C). Cells were harvested 48 hr post - transfection and solubilized in TBS (20 mM Tris pH 8.0 at 4°C, 150 mM NaCl) with 10 mM MNG (Antrance), 0.1% CHS (Antrance) and protease inhibitors including 1 μg/ml aprotinin, 1 μg/ml leupeptin and 1 μg/ml pepstatin on ice for 30 min. Cell lysates were cleared by centrifuge at 40,000 rpm for 30 min and supernatants were loaded onto Superose 6 increase (GE Healthcare) for FSEC analysis ^51^.

### FLIPR membrane potential assay

hTRPC6 constructs were transfected into AD 293 cells (grown in DMEM+10% FBS, 37°C) and seeded on 384-well plates at the density of ∼10000 cells/well for FLIPR assay using FLIPR Membrane Potential Assay Reagent (Thermo Fisher). Fluorescence signals were read with excitation/emission at 510-545/565-625 nm. EC_50_ was obtained by fitting the data into the function: Y = Bottom + (Top-Bottom)/(1+10^(LogEC_50_-X)) and IC_50_ was obtained by fitting the data into function: Y = Bottom + (Top-Bottom)/(1+10^(X-LogIC_50_)).

### Protein expression and purification

We established the BacMam expression system for large scale expression of full length hTRPC3 and hTRPC6 as describe before ^50^. Briefly, full length hTRPC6 was subcloned into a home-made BacMam vector with N terminal strep-GFP tag. Bacmid was then generated by transforming this construct into DH10Bac *Escherichia coli* cells. Baculovirus was harvested about 1 week after transfecting bacmid into Sf9 cells cultured in Sf-900 III SFM medium (Gibco) at 27°C.When Free Style 293-F cells (grown in free style 293 medium + 1% FBS, 37°C) in suspension were grown to a density of 2.0 × 10^6^ ml^−1^,baculovirus was added to it. 12 hr after infection, 10 mM sodium butyrate and 100 nM BTDM were added and temperature was lowered to 30°C. Cells were harvested 60 hr post-infection and washed twice using TBS buffer (20 mM Tris pH 8.0 at 4°C, 150 mM NaCl). Cell pellets were frozen at −80°C for use.

Cell pellets were resuspended in TBS buffer containing 10 mM MNG, 0.1% CHS, 100 nM BTDM and protease inhibitors including 1 μg/ml aprotinin, 1 μg/ml leupeptin, 1 μg/ml pepstatin, 1 mM PMSF and rotated at 4°C for 1 h. Unbroken cells and large cell debris were removed by centrifugation at 14,800 rpm for 10 min. Supernatants were further centrifuged at 40,000 rpm for 40 min in Ti45 rotor (Beckman). After centrifugation, appropriate amount of purified GST-tagged GFP-nanobody ^52^ was added to the supernatant and rotated at 4°C for 10 min. Then samples were loaded on 7 ml GS4B resin (GE Healthcare). Resin was washed with wash buffer 1 (TBS + 40 μM GDN + 0.01 mg/ml soybean lipids + 100 nM BTDM). Then it was further washed with wash buffer 2 (wash buffer 1+ 10 mM MgCl_2_ + 1 mM ATP) to remove bound chaperones. Protein was eluted with elution buffer (10 mM reduced glutathione + 50 mM Tris (pH 8.0 at RT) + 40 μM GDN + 0.01 mg/ml soybean lipids + 100 nM BTDM + 4 mM DTT) and concentrated using 100-kDa cut-off concentrator (Millipore). To cleave the tags off the hTRPC6 protein and remove glycosylation, concentrated proteins were incubated with H3CV protease and PNGase F at 4°C overnight. Proteins were further purified by Superose 6 increase (GE Healthcare) running in buffer containing TBS, 40 μM GDN, 0.01 mg/ml soybean lipids, 100 nM BTDM and 1 mM TCEP. The peak fractions corresponding to tetrameric TRPC channel were collected, concentrated and mixed with soybean lipids, purified MSP2N2 (prepared as described previously ^53^) at a molar ratio of TRPC6 monomer: MSP2N2: soybean lipids= 1: 7: 225 to initiate nanodisc reconstitution. After incubating the mixture at 4°C for 30 min, Bio-beads SM2 (Bio-Rad) were added and the reconstitution system was rotated at 4°C for 1 h before another batch of fresh bio-beads were added. The system was rotated at 4°C overnight. Bio-beads were removed and the reconstitution system were cleared by centrifugation at 40,000 rpm for 30 min before loaded onto a Superose-6 increase column running in TBS containing 1 mM TCEP. The peak fractions corresponding to tetrameric hTRPC6 channel in nanodiscs were assessed by SDS-PAGE.

### EM sample preparation

Purified hTRPC6 nanodisc was concentrated to A280 = 4.5 with estimated concentration of 35.8 μM monomers and supplemented with 100 μM BTDM. To overcome the preferred orientation of particles in ice, 0.5 mM non-solubilizing detergent fluorinated octyl-maltoside (FOM) ^54^ was added to the sample before cyro-EM sample preparation. Protein sample were loaded onto glow-discharged GiG R1/1 holey carbon grids and plunged into liquid ethane by Vitrobot Mark IV (Thermo Fisher) at Institute of Biophysics, Chinese Academy of Sciences.

### Data Collection

The cryo-grids were firstly screened on a Talos (Thermo Fisher) operated at a voltage of 200 kV with an eagle 4k × 4k CCD camera (Thermo Fisher). For hTRPC6 data collection, grids were imaged with a GatanK2 Summit direct detector camera on Titan Krios (Thermo Fisher) at the voltage of 300 kV, with a Cs corrector, magnification of 37,500 ×, equivalently to the pixel size of 1.33 Å, and with defocus values ranging from −1.0 µm to −3.0 µm. Super-resolution movies (50 frames per movie) acquisition were performed automatically using Serial EM ^55^ with a dose rate of 8 e^−^/pixel/s on detector and a total dose 50 e^−^/Å^2^.

### Image processing

For hTRPC6, 1,868 movies were collected. MotionCor2 ^56^ was used for image binning, motion correction and dose weighting. After motion correction, about 120 bad micrographs were removed. GCTF ^57^ was used for CTF estimation of all the motion-corrected sums (1,748 movies). After manual picking particle, reference-free 2D classification was performed using Relion2.0 ^58^. The resulting 2D averages were used as templates for autopicking on all micrographs. Repeated 2D classification was used to remove bad particles, leaving 189,781 particles for 3D classification. An *ab initio* 3D reconstruction was generated using cryoSPARC ^59^. This initial model was used as the initial reference for 3D classification in Relion2.0 without any symmetry restrains. After 3D classification, one class containing 53,528 particles with clearly visible α helices was selected for 3D refinements in cryoSPARC using C4 symmetry. The reconstruction reaches 3.8 Å, based on gold standard FSC 0.143 cut-off after correction of masking effect ^60^. The map was sharpened with a B-factor of −166.8 Å^2^ determined by cryoSPARC automatically.

### Model building

*Ab initio* model of hTRPC6 was built using EM builder ^59^ and Rosetta CM ^61^. The model was further manually rebuilt using Coot ^62^, using the structure of NOMPC (5VKQ) as reference of TMD. Structure was refined with PHENIX ^63^ against sharpened map. Pore radius was calculated with HOLE ^64^.

### Quantification and statistical analysis

The orientation distribution of the particles used in the final reconstruction was calculated using RELION 2.0 ^66^. The local resolution map was calculated using ResMap ^67^.

### Data availability

3D cryo-EM density map of hTRPC6 has been deposited to the Electron Microscopy Data Bank (EMDB) under the accession number: EMD-XXXX. Coordinates of atomic model of hTRPC6 has been deposited in the Protein Data Bank (PDB) under the accession number: XXXX. All other data are available from the corresponding authors upon reasonable request.

## Acknowledgements

We thank all of Chen Lab members for kindly help. Cryo-EM data collection was supported by the National Center for Protein Science (Shanghai) with assistance of Liangliang Kong and Zhenglin Fu, Electron microscopy laboratory and Cryo-EM platform of Peking University with the assistance of Xuemei Li, and Center for Biological Imaging, Institute of Biophysics, Chinese Academy of Science with assistance of Zhenxi Guo. Part of structural computation was also performed on the Computing Platform of the Center for Life Science and High-performance Computing Platform of Peking University. The work is supported by grants from the Ministry of Science and Technology of China (National Key R&D Program of China, 2016YFA0502004 to L. C.), National Natural Science Foundation of China (31622021 and 31521062 to L. C.), Young Thousand Talents Program of China to L. C. and the China Postdoctoral Science Foundation (2016M600856 and 2017T100014 to J.-X.W.). J.-X. W. is supported by the postdoctoral foundation of the Peking-Tsinghua Center for Life Sciences, Peking University.

## Author contributions

X. Z. and L. C. initiated the project. Q. T., W. G., L. Z., M. L., X. Zhou, X. Z., and L. C. designed the experiments. Q. T., W. G., J.-X. W., and L. C. prepared the EM sample, collected the EM data, performed image processing and built the model. Q. T. and W. G. generated mutants. L. Z., M. L., X. Zhou, and X. Z. provided BTDM, performed electrophysiology recording and FLIPR assay. Q. T., W. G., J.-X. W., and L. C. wrote the manuscript draft. All authors contributed to manuscript preparation.

## Supplementary information

Supplemental Information includes seven figures, two tables and one movie.

**Figure S1.**
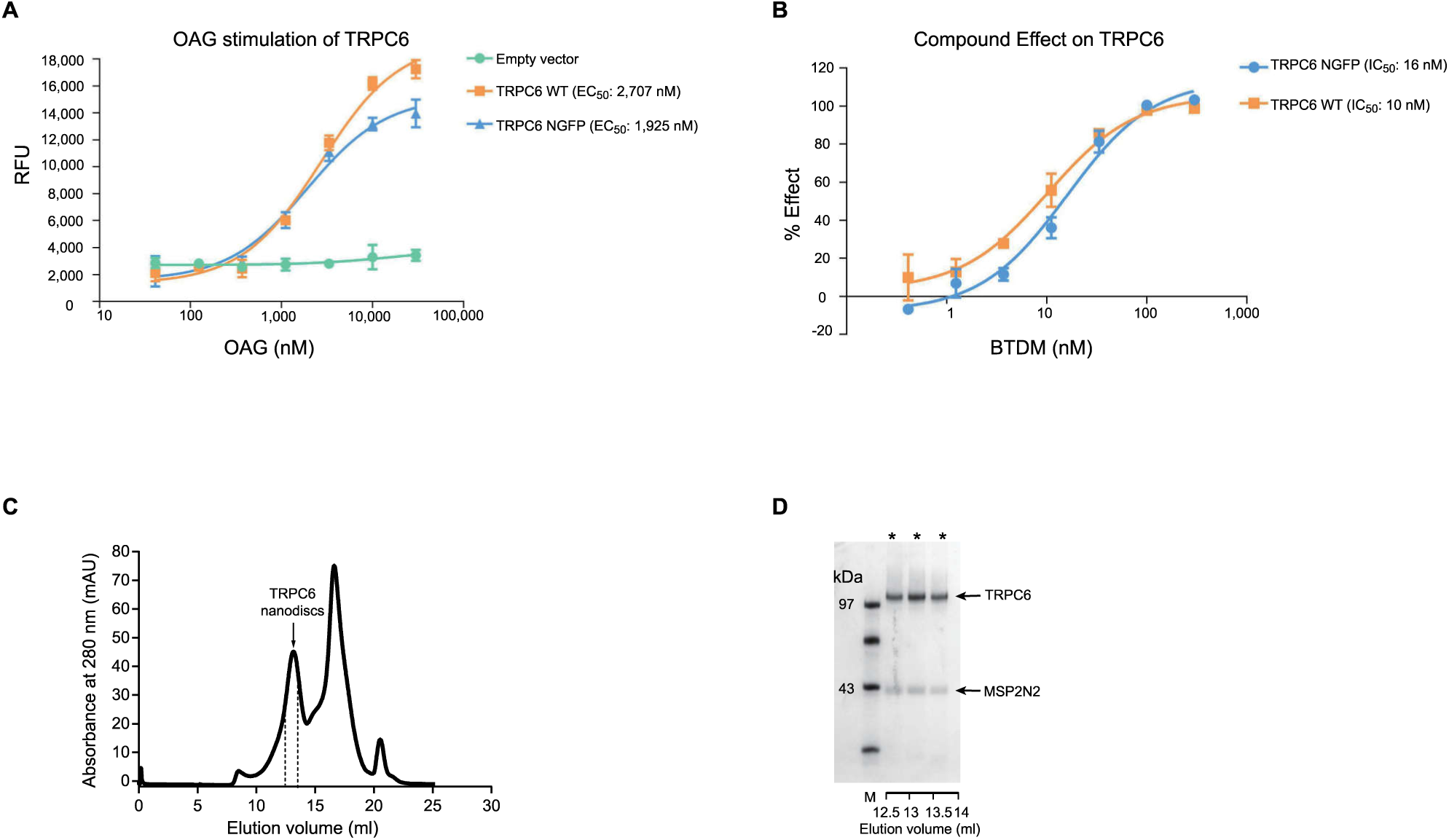
Functional characterization and purification of NGFP-hTRPC6. **(A)** Activation effect of OAG on NGFP-hTRPC6 constructs, measured by FLIPR membrane potential assay (n=3). **(B)** Inhibition effect of the BTDM on NGFP-hTRPC6 constructs (n=3). **(C)** Representative size-exclusion chromatography of hTRPC6 reconstituted into nanodiscs. Fractions between dashed lines contain hTRPC6 in nanodiscs. **(D)** SDS-PAGE of size-exclusion chromatography peak fractions. Fractions denoted by asterisks were pooled for cryo-EM analysis.

**Figure S2.**
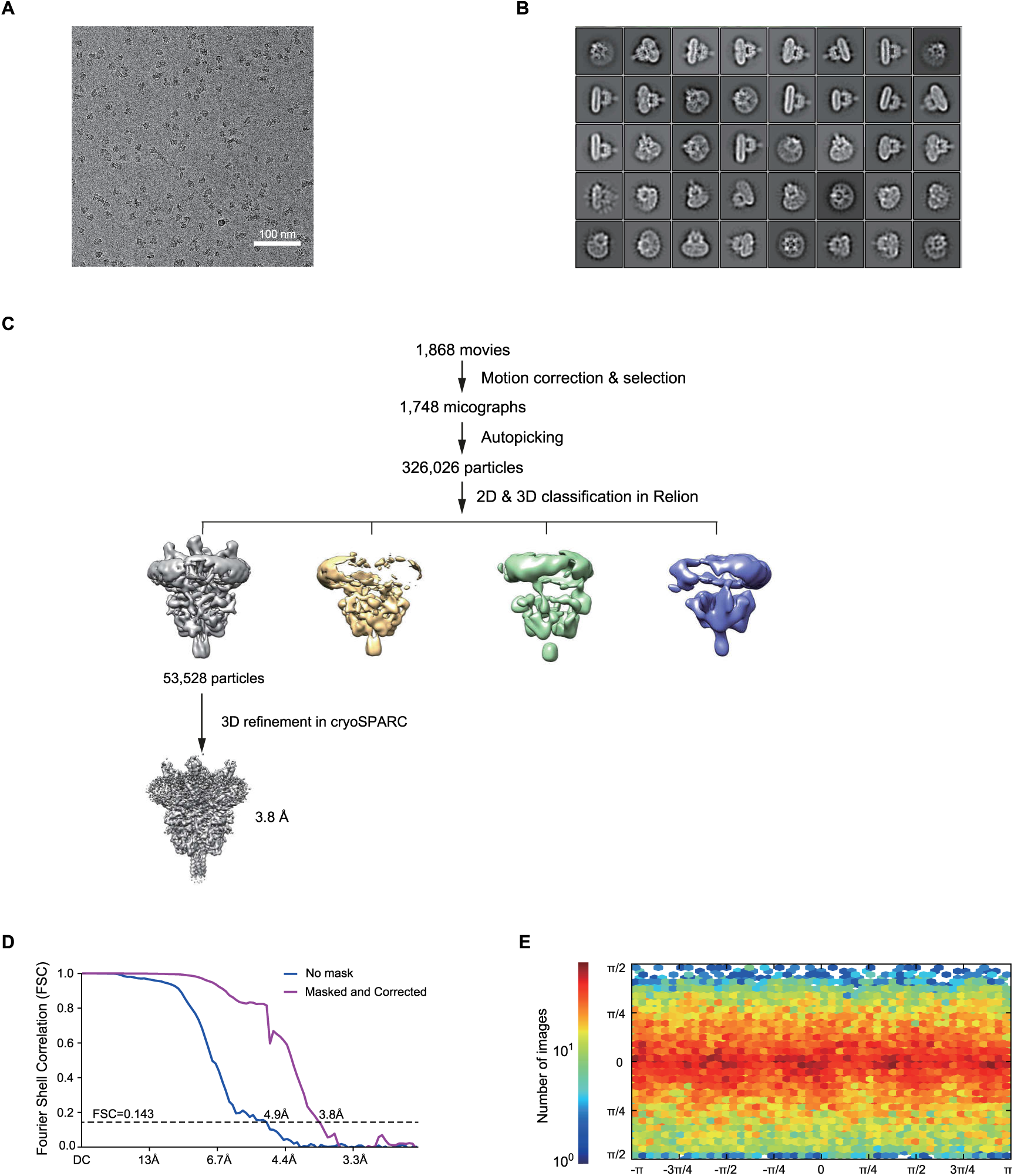
Cryo-EM image analysis of the hTRPC6 channel in nanodiscs. **(A)** Representative raw micrograph of hTRPC6 nanodisc particles recorded on K2 Summit camera at a defocus of −2.0 µm. **(B)** Representative 2D class averages of hTRPC6 in nanodiscs. **(C)** Flowchart of hTRPC6 cyro-EM data processing. **(D)** FSC curves of the two independently refined maps for unmasked (blue line, 4.9 Å) and masked & corrected (purple line, 3.8 Å). Resolution estimation was based on the criterion of FSC 0.143 cut-off. **(E)** Angular distribution histogram of the final hTRPC6 reconstruction. This is a standard output from cryoSPARC.

**Figure S3.**
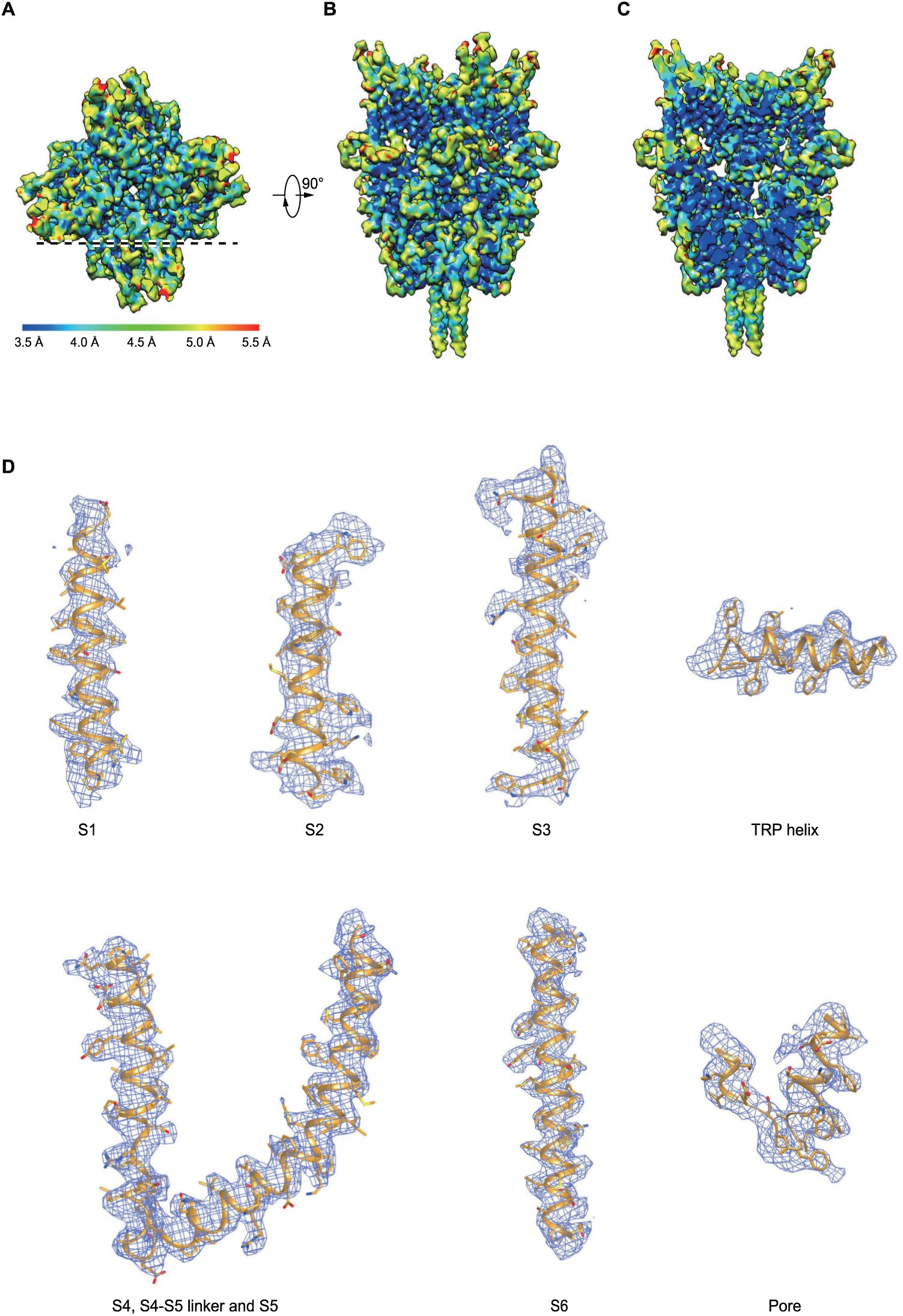
Cryo-EM map of hTRPC6 in nanodiscs. **(A-C)** Top view **(A)**, side view **(B)** and cross-section **(C)** of the final sharpened hTRPC6 cryo-EM density map colored by local resolution estimated by Resmap. A dashed line in **(A)** indicates the position of cross-section of **(C)**. **(D)** Cryo-EM density map of representative transmembrane helices. The density maps in light blue mesh are superimposed with the hTRPC6 atomic model in orange.

**Figure S4.**
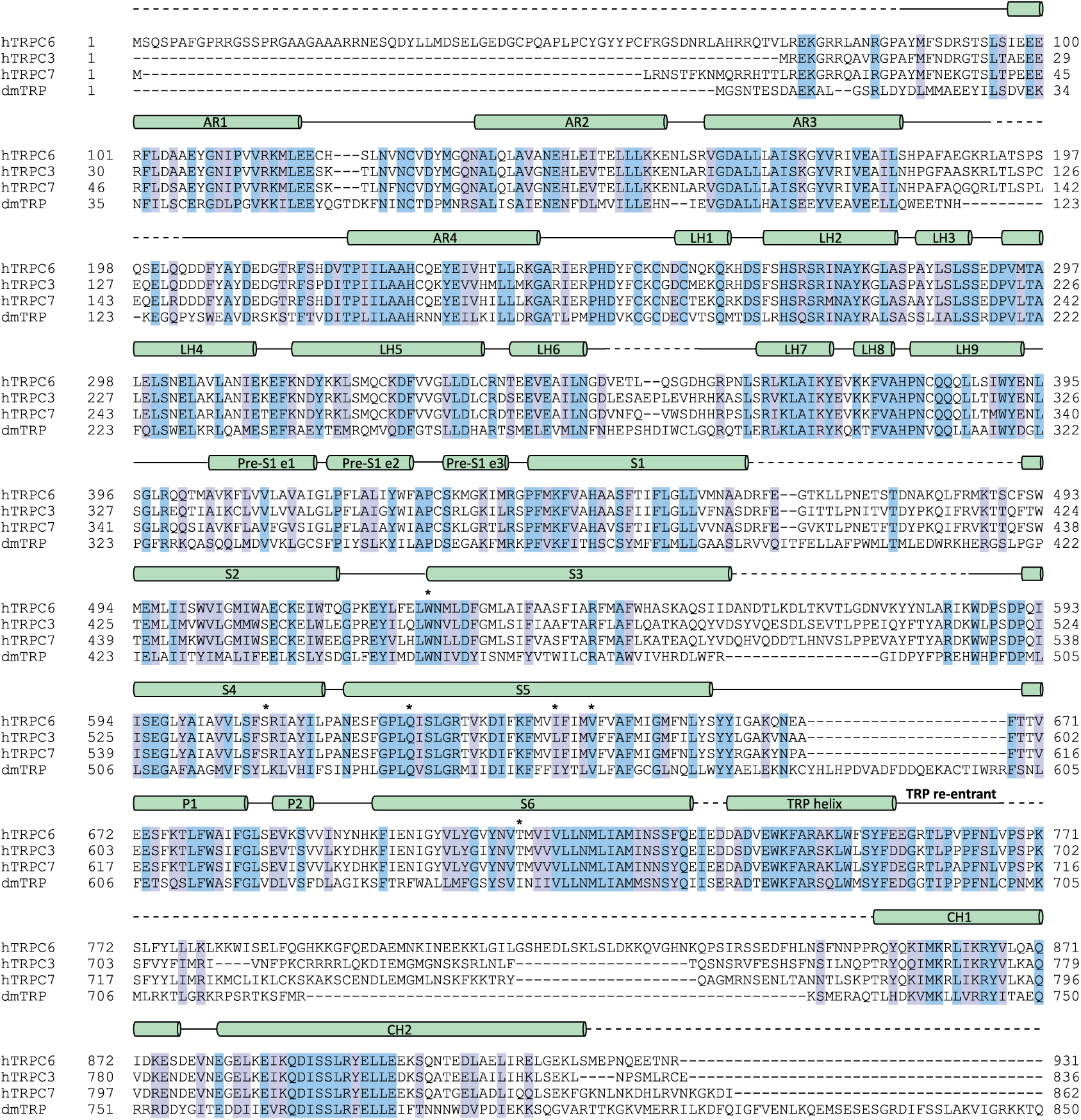
Sequence alignment of humanTRPC6 (hTRPC6), hTRPC3, hTRPC7 and Drosophila TRP (dmTRP). Secondary structure elements are depicted above sequences (cylinders for α helices, lines for loops and dashed lines for unmodeled residues). Conserved residues are highlighted in purple and highly conserved residues are in blue. The residues that are involved in BTDM binding are labeled with asterisks. The non-conserved extended C-terminus of Drosophila TRP channel is not shown due to limited space.

**Figure S5.**
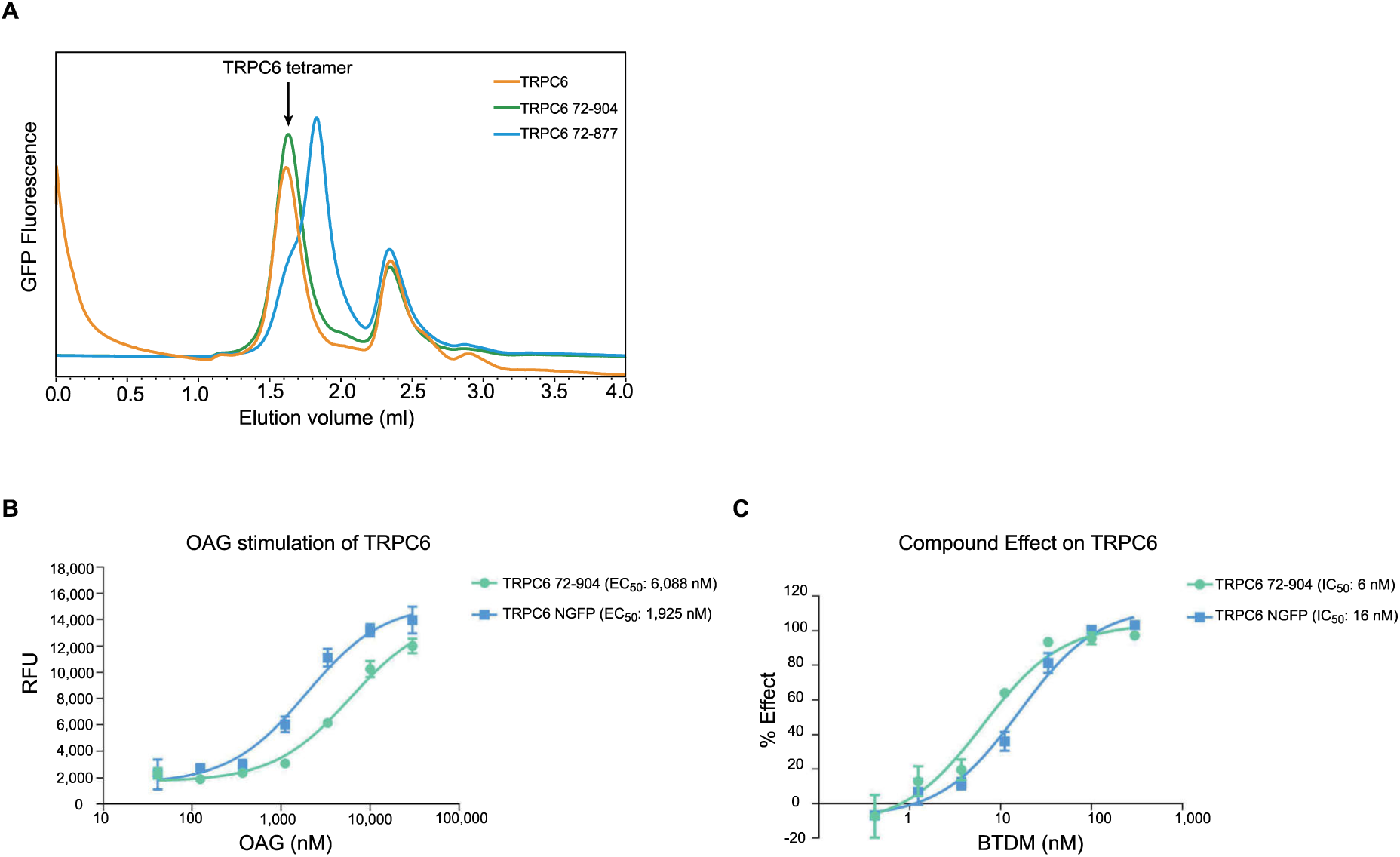
Characterization of hTRPC6 truncation constructs. **(A)** FSEC profile of hTRPC6 truncation constructs. Arrow indicates the peak position corresponding to the tetrameric N-terminal GFP-tagged hTRPC6. **(B)** Activation effect of OAG on hTRPC6 72-904 construct, measured by FLIPR membrane potential assay (n=3). **(C)** Inhibition effect of the BTDM on hTRPC6 72-904 construct (n=3).

**Figure S6.**
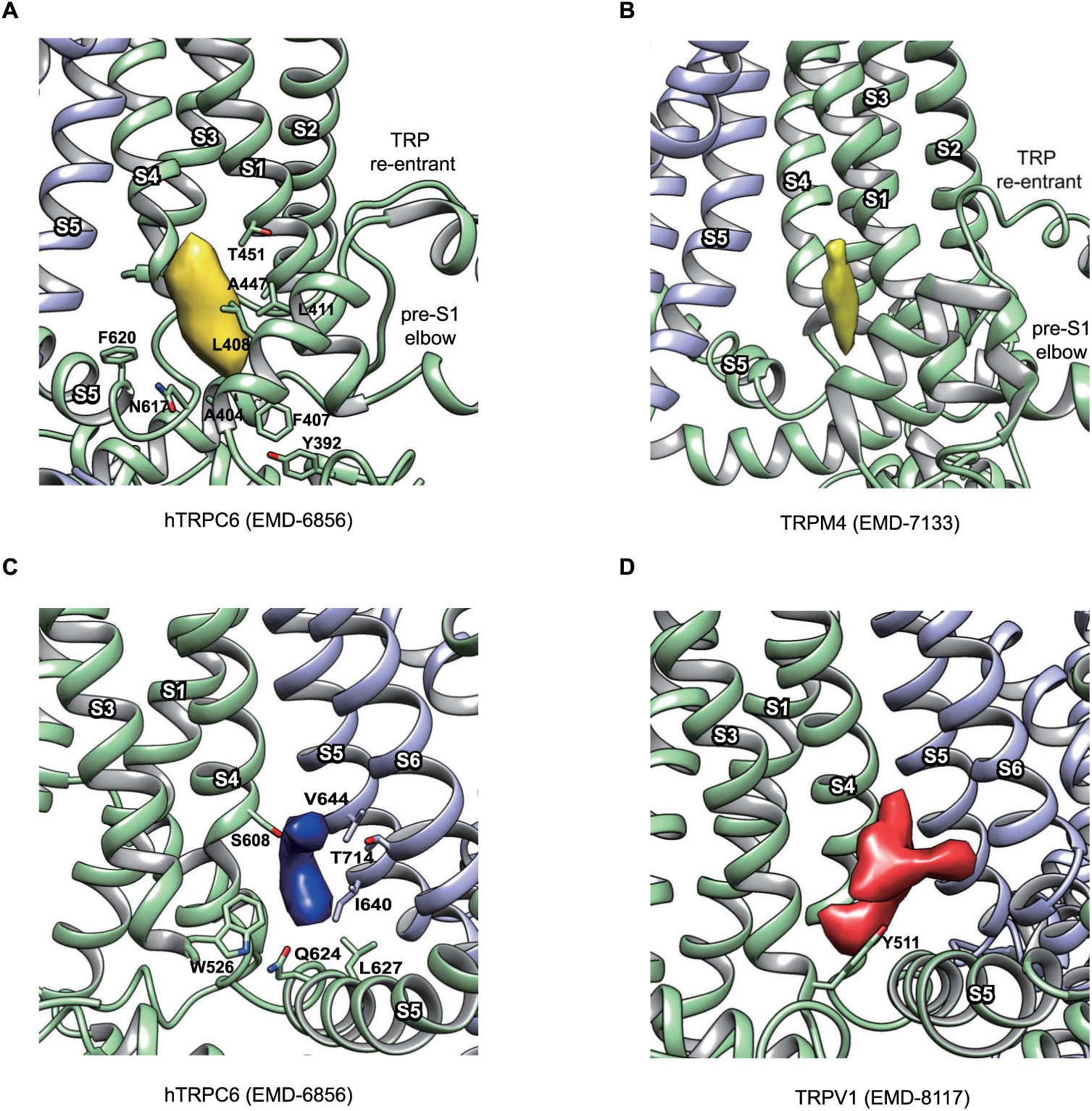
Ligand densities in the cryo-EM map of hTRPC6. **(A)** The putative CHS density in hTRPC6 cryo-EM map is shown in yellow. Residues (Y392, A404, F407, L408, L411, A447, T451, N617, and F620) close to the density are shown as sticks. **(B)** The CHS density of TRPM4 (EMD-7133) shown in yellow is at the similar position to hTRPC6. **(C)** The BTDM density is colored in dark blue. The side chain of W526, S608, Q624, L627, I640, V644, and T714 close to the density are shown as sticks. **(D)** The density of TRPV1 activator RTX (EMD-8117) shown in red is at the similar position to BTDM of TRPC6 in panel **(C)**.

**Figure S7.**
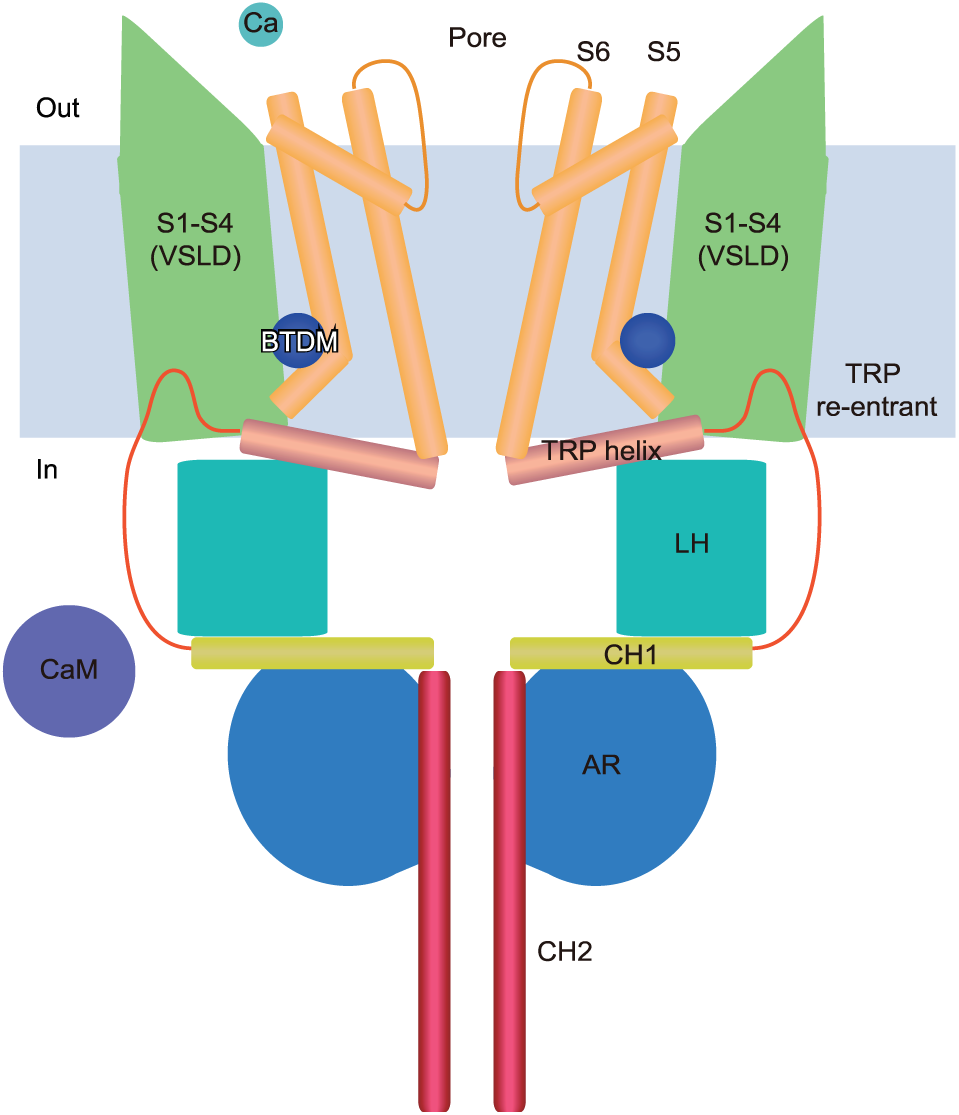
Model for TRPC channel regulation and inhibition. Side view of the cartoon model of TRPC channel in apo state. TRPC is colored the same as in Figure 2A. For simplicity, only two TRPC subunits are shown. The extracellular calcium ion (Ca), intracellular calmodulin (CaM) and BTDM are shown as cyan, purple and blue spheres, respectively.

**Table S1:** EC_50_ of OAG and IC_50_ of Inhibitor on hTRPC6 constructs.

| TRPC6 constructs | OAG |  | Inhibitor |  |
| --- | --- | --- | --- | --- |
|  | LogEC<sub>50</sub><sup>a</sup> | EC<sub>50</sub> (nM) | LogIC<sub>50</sub><sup>a</sup> | IC<sub>50</sub> (nM) |
| TRPC6 WT | 3.43 ±0.06 | 2,707 | 1.01 ±0.07 | 10 |
| TRPC6 NGFP | 3.28 ±0.08 | 1,925 | 1.20 ±0.07 | 16 |
| **Mutations in CHS** |  |  |  |  |
| **binding pocket** |  |  |  |  |
| Y392F | 3.32 ±0.09 | 2,100 | 1.26 ±0.11 | 18 |
| A404C | 3.41 ±0.07 | 2,589 | 1.30 ±0.10 | 20 |
| F407A | 3.25 ±0.06 | 1,774 | 1.13 ±0.07 | 13 |
| L408M | 3.38 ±0.09 | 2,377 | 1.18 ±0.07 | 15 |
| L411F | 3.40 ±0.14 | 2,494 | 1.35 ±0.11 | 23 |
| A447T | 4.02 ±0.08 | 10,481 | 1.07 ±0.08 | 12 |
| A447F | 3.25 ±0.05 | 1,779 | 1.76 ±0.13 | 57 |
| T451F | 3.75 ±0.09 | 5,684 | 1.19 ±0.09 | 16 |
| T451L | 3.29 ±0.05 | 1,944 | 1.35 ±0.13 | 22 |
| N617S | 3.81 ±0.10 | 6,406 | 0.6 ±0.08 | 4 |
| F620L | 3.09 ±0.11 | 1,227 | 0.79 ±0.08 | 6 |
| **Mutations in BTDM** |  |  |  |  |
| **binding pocket** |  |  |  |  |
| W526A | 2.93 ±0.06 | 850 | 2.00 ±0.14 | 100 |
| S608L | 2.51 ±0.07 | 326 | - | >10,000<sup>b</sup> |
| Q624A | 3.70 ±0.07 | 4,990 | 3.39 ±0.16 | 2,463 |
| I640F | 4.07 ±0.13 | 11,709 | 1.62 ±0.14 | 42 |
| V644F | 2.93 ±0.03 | 857 | - | >10,000<sup>b</sup> |
| T714F | 3.17 ±0.10 | 1,463 | 3.93 ±0.17 | 8,430 |
| **Gain of function mutants** |  |  |  |  |
| P112Q | 3.32 ±0.69 | 2,090 | 1.15 ±0.07 | 14 |
| R895C | 3.27 ±0.09 | 1,852 | 1.22 ±0.09 | 17 |
| E897K | 3.48 ±0.08 | 2,993 | 0.97 ±0.12 | 9 |

<sup>a</sup>Data were expressed as logEC<sub>50</sub> ( or logIC<sub>50</sub>) ±SEM, n=3.<sup>b</sup>The highest inhibitor concentration (10,000 nM) inhibited the activation of TRPC6 for less than 30%.

**Table S2:**
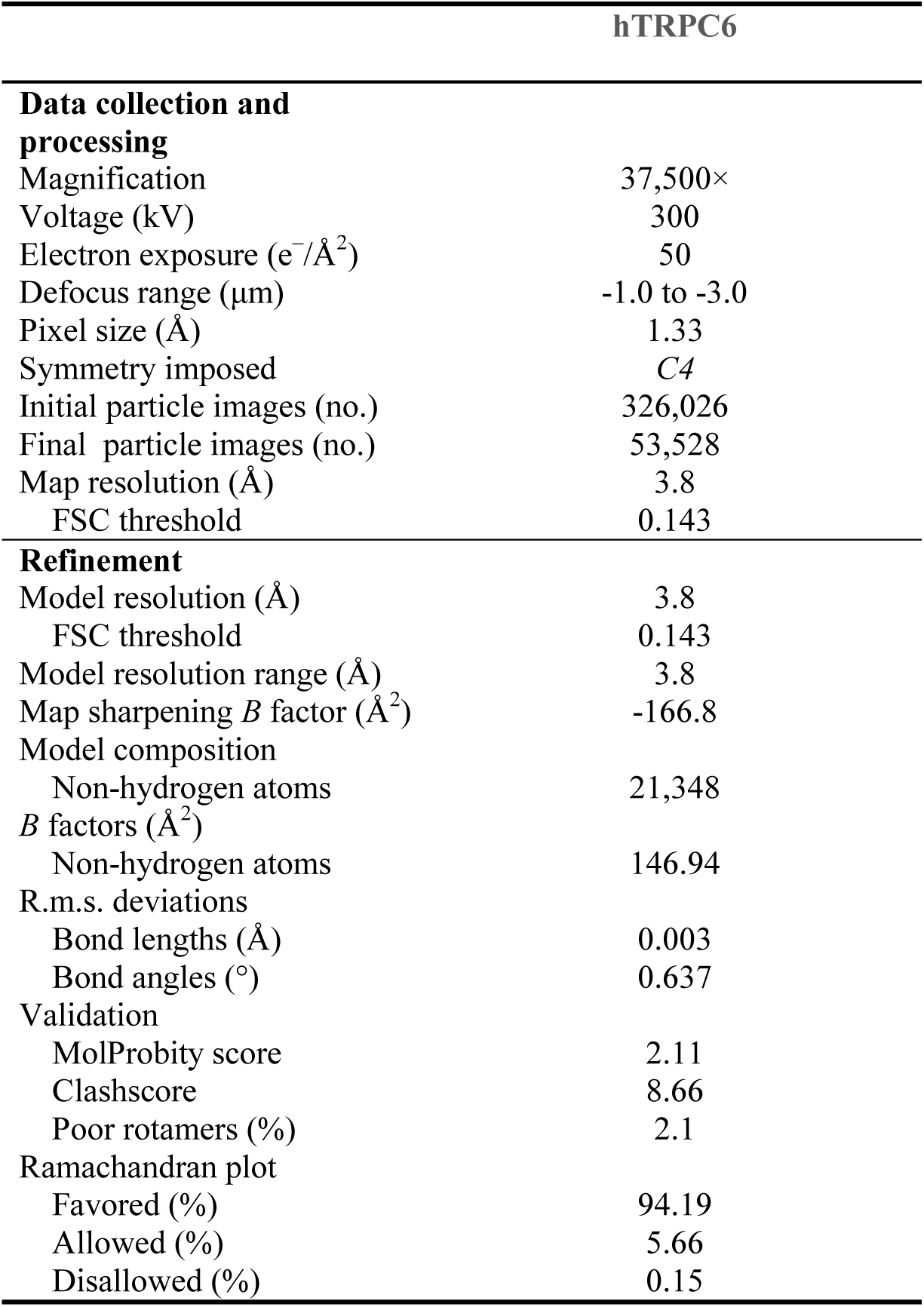
Cryo-EM data collection, refinement and validation statistics.

